# Comparative Analysis of Cell Ultrastructure with Evidences of Programmed Death in Tissue Cultures of Cereals

**DOI:** 10.1101/186676

**Authors:** Bishimbayeva N.K., Rakhimova E.V.

## Abstract

Degrading cells of wheat and barley callus have all known ultrastructural features, which are characteristic for apoptotic cells: intense vacuolization, increasing the size of the nucleus, cupped invaginations of plastids, mitochondria saving until the last stages of cell degradation and the formation of vesicles, similar to apoptotic bodies. It is the first showed that degrading cells in embryogenic callus are characterized with the destruction of the nuclear membrane and the allocated substances of fibrillar nature in the periplasmic space and on the surface of cells.

---

Programmed cell death (PCD, apoptosis, apoptotic-like programmed cell death) is a process of controlled degradation and destruction of cells, which plays an important role in the ontogeny of plants and occurs in plants during development. It is involved in the formation of female gametes, male sexual organs and seed storage tissues in plants. Programmed cell death also occurs during the embryogenesis and germination of plants (Palavan-Unsal et al., 2005), xylogenesis, parenchyma formation, several plant reproductive processes, seed development ((Pennel, Lamb, 1997; Gray, 2004). Apoptosis is typical for leaf aging and dying (Noodén et al., 1997). In addition, programmed cell death is well documented in relation to manifestation of hypersensitive response caused by the interaction between the host plant and an incompatible pathogen (Hatsugai et al., 2004).

According to classification of some authors (Van Doorn et al., 2011) based on morphological criteria, the use of the term ‘apoptosis’ is not justified in plants, but at least two classes of programmed cell death can be distinguished: vacuolar cell death and necrosis. Plant vacuolar cell death also is described as autophagy and classification of autophagy to several types was given by some authors (Van Doorn et al., 2013). Critical analysis of considerable number of articles describing ‘plant apoptosis’ or ‘apoptotic-like programmed cell death (PCD)’ reveals three major points that indicate misuse of the term ‘apoptosis’, the most important of which: chromatin condensation are often quoted as apoptotic features. However, this morphological feature is not specific to apoptosis, and can be observed during necrosis and autophagic death (Lee et al., 2001; Hoyer-Hansen et al., 2005; Fink, Cookson, 2005). At the same time, other authors (Jan et al., 2008) note that morphological and biochemical features of apoptosis appear to be similar across the eukaryotic kingdom, this suggests that despite differences, there may be a functional similarity between plant and animal apoptosis.

The most common ultrastructural features of programmed cell death observed are: reduction of cellular volume, formation of periplasmic space, vacuolization and fragmentation of cytoplasm into small parcels called apoptotic bodies, formation of huge vacuoles in the cells (Kirnos et al., 1997; Groover, Jones, 1999; Drew et al., 2000; Zamyatnina et al., 2002), appearance in the vacuole of vesicles containing portions of the cytoplasm with active organelles, subsequent destruction of tonoplast, chromatin condensation in the nucleus (O’Brien et al., 1998), nuclear segmentation and subsequent destruction of the nucleus (Kirnos et al., 1997; Bakeeva et al., 2001), and very little ultrastructural modification of cytoplasmic organelles. During programmed cell death apoptotic cells in plants, because of their solid cell walls, do not disappear completely. They take part in formation of vascular bundles and aerenchyma (Drew et al., 2000).

Ultrastructural aspects of apoptosis, including condensation and fragmentation of the nucleus, cell shrinkage, membrane blebbing, as well as the formation of apoptotic bodies were first observed by Kerr and co-workers (Kerr et al., 1972). Apoptotic bodies have not been shown to be present during plant cell death. This is due to the absence of phagocytosis and presence of the cell wall (Jan et al., 2008). Spherical fragments of protoplasts (apoptotic-like bodies) of similar appearance were observed in aging cell cultures (Groover et al., 1997), the contents of a plant cell, packaged in such apoptotic-like bodies, release from a dying cell into tissue (Wang et al., 1996).

Many authors (Samuilov, 2001; Vanyushin et al., 2004 and others) have noted that data on apoptosis in plants are still fragmentary, and apoptosis in plants has many specific features that are not observed in animals.

In this paper we have tried to show ultrastructural features of programmed death of cells detected earlier (Bishimbayeva, 2006; Bishimbayeva et al., 2007a, b) in the embryogenic callus of cereals.

## Materials and methods

The objects of study were embryogenic calluses of barley and wheat. Long-cultivated friable embryogenic calluses were derived from immature embryos (1.0-1.4 mm), isolated from the milky-wax ripeness seeds of barley and spring wheat cultivars (Arna and Otan) milky-wax ripeness. Barley calluses were cultured on an agar medium Gamborg B5 (Gamborg, Eveleigh, 1968) supplemented with 7.0 mg/l 2,4-D, wheat calluses - on Murashige and Skoog medium (Murashige, Skoog, 1962) supplemented with 5.0 mg/l 2,4-D.

For the electron microscopy calluses were transferred from the surface of artificial medium into a fixing solution, prefixed for 2.5 h at room temperature in 2% glutaraldehyde buffered with sodium cacodylate, washed four times with freshly prepared same buffer, and postfixed in 1 % buffered osmium tetroxide for 2 h at room temperature. Specimens were stained with 2 % alcoholic uranyl acetate for 2 h, dehydrated in graded series of ethanol and 100 % acetone, embedded in Epon-Araldite, and incubated for 48 h at 60°C. Ultrathin sections were cut with glass knives using a Reichert Ultracut Ultratome (Austria), collected on formvare coated copper grids, and analyzed with Jem-100B transmission electron microscope operated at 80 kV.

The size of the nuclei and cell organelles was determined by electron micrographs of median sections.

## Results and discussion

### Cell wall and plasmalemma (plasma membrane)

The cell walls of wheat calluses and barley have moderate electron density; sometimes modifications represented invaginations – deep ingrowths into the cell are observed (fig. 1a-c). The electron density of these invaginations is low and corresponds to the density of young cell walls. However, deposition as bubbles and moderate electron-dense lumps may be observed in electron translucent material.

**Figure 1.**
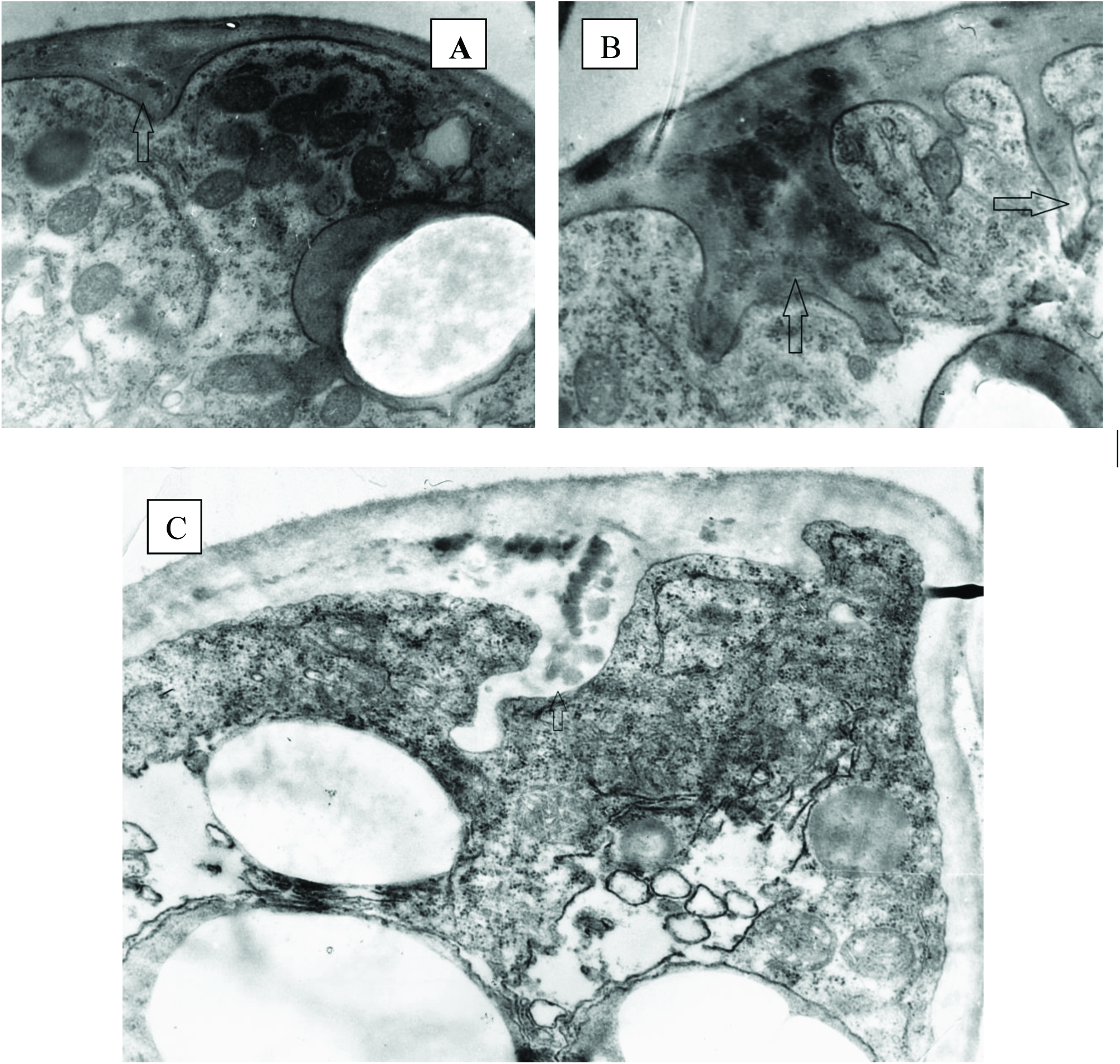
Modifications of cell walls (arrows) of barley (A, B) and wheat (C) calluses.

It seems that such modifications of the cell wall are either a reaction to the composition of the culture medium, or mechanism of cytokinesis changes (perhaps - is blocked or broken) under the influence of the medium, and the formation of the cell wall is irregular. Some authors (Vanyushin, 2001) observing similar defective changes of cell walls, believe that the invaginations are the result of incomplete cytokinesis, which causes formation of partial cell wall. Perhaps the modifications of the cell walls are connected with the process of fragmentation of the cytoplasm (Zamyatnina et al., 2002).

In the final stages of programmed cell death of wheat and barley callus plasmalemma integrity is most often maintained, but there invaginations and lomasomes appear. Similar invaginations of plasma membrane are also characteristic for vacuolated callus cells of *Hibiscus rosa-sinensis* (Li et al., 1998) and callus tissue derived from embryogenic structures of wheat (Kosulina, Lapikova, 1983). In some cells there is separation of plasma membrane from the cell wall – pseudo-plasmolysis. This phenomenon is considered as the best evidence of apoptosis and was observed in plant cells by other researchers (O’Brien et al., 1998; McCabe et al., 1997; Gao, Showalter, 1999).

Presence of vesicles and dense globules in the periplasmic space is frequently observed during pseudo-plasmolysis in barley callus cells.

### Nucleus

The nucleus of wheat callus cells is quite large, with an average of (10.9 x 8.9) μm, the central position of nucleus is rare, and it is likely to be associated with progressive cell vacuolization. On sections the nuclear envelope is slightly wavy; there are rare ribosomes on the outer membrane. Type of nucleus is chromonemic, which is typical for a physiologically active cell. Condensed chromatin is presented with long, highly-contrasting strands – chromonemata. Some chromonemata contact with the nuclear envelope.

In the initial stages of apoptosis the nucleus does not change, but chromonemata become more contrasting. In some nuclei distinct changes are observed in chromatin morphology, such as chromatin condensation and margination at the nuclear envelope. Similar changes in morphology of chromatin are observed in the nuclei of apoptotic cells in the apical part of the initial leaf (in 5 day old) etiolated wheat seedlings and in wheat coleoptile (in the nuclei of cells with vacuolar vesicles) (Zamyatnina et al., 2002). Condensation and margination of chromatin are well known specific ultrastructural features of the initial stages of apoptosis in both animals and plants (Vanyushin, 2001; Vanyushin et al., 2004). The nucleus often looks intact even in case of partial destruction of the nuclear membrane and in case of the total destruction of the nuclear membrane condensed chromatin is retained in the cytoplasm. These phenomena in the early stage of cellular injury are thought to result in nuclear degradation and wheat cell death in the later stage.

The nucleus of barley callus cells is quite large, with an average of (9.9 x 4.6) μm. Despite intensive ongoing processes of autolysis during apoptosis cells retain an active nucleus.

### Nucleolus

Prominent nucleolus of wheat cells, an average of (3.6 x 2.5) μm, has nucleolar vacuoles, whose presence is considered to be one of the features of the secretory cells (Shubnikova, 1967). In some nuclei two nucleoli are observed, one of which is significantly larger than the second (2.6 x 2.2) μm and (1.8 x 1.53) μm, respectively). It is interesting to note that in the process of apoptosis nucleolus may disappear or remain as a long time. These nucleoli become larger and their number reaches three. The nucleoli appear intact even in case of the partial destruction of the nuclear membrane.

Two active nucleoli, (2.7 x 1.6) μm, with nucleolar vacuoles are often observed in barley callus cells. During the apoptosis nucleoli are stored for a long time.

### Ribosomes

In barley and wheat callus cells ribosomes occur in significant number. They can be free, or associated with membranes of endoplasmic reticulum - on its outer membrane surface. Ribosomes are also represented by polyribosomal complexes. Ribosomes can be observed on the outer surface of the membrane of vesicles, concentrated in the cell walls of wheat and barley callus (the so-called “fringed vesicles”).

### Golgi apparatus

In barley and wheat callus cells the Golgi apparatus consists of 3-12 cisternae, from which the vesicles are separated. It should be noted that among these vesicles are not observed “fringed vesicles” with ribosomes.

### Endoplasmic reticulum

The endoplasmic reticulum of wheat callus cells is well developed. Cisternae of endoplasmic reticulum are located mainly around the nucleus, forming a so-called “reticular case” and on the periphery of the cell, parallel to the cell wall, contacting the plasma membrane. In some cells short tubules and cisternae of endoplasmic reticulum form a dense extensive network. The ratio of the smooth and the granular elements of endoplasmic reticulum varies greatly from cell to cell.

It is known that the rough endoplasmic reticulum is involved in the synthesis of glycoproteins, smooth reticulum - in the synthesis of terpenoids and transport of substances within the cell (Vassilyev, 1977).

The endoplasmic reticulum of barley callus cells form a dense network of short curved tubules. The longest cisternae of endoplasmic reticulum are characteristic for the cell periphery. Short cisternae often surround vacuoles and mitochondria, sometimes forming contacts. Despite intensive ongoing processes of apoptosis cells keep their well-developed endoplasmic reticulum, which is a feature of metabolic activity. Presence of contacts between endoplasmic reticulum, mitochondria and nuclei, is characteristic for the callus cells of barley and noted in cells with strong vacuolization. In the callus cells, where pseudoplasmolysis is noted, cisternae of endoplasmic reticulum break.

### Plastids

Plastids of wheat callus cells are characterized by high variability of their shape. Most of them are elongated (3,2 x 1,2) μm, but there are as dumbbell, and organelles with irregularly shaped. Double membrane envelope is not expressed in some of the plastids, and there is only one membrane. Prolamellar bodies, the initials of the lamellar system or 1-2 small plastoglobules are marked in others plastids. Similar features of plastids (absence of complete envelope, underdeveloped lamellar system and irregular shape of organelle) are also characteristic for chlorophyll-deficient mutants of *Festuca pratensis* (Venzhik et al., 2002) and albino regenerates of spring wheat (Galiyeva, 2001; Galiyeva et al., 2001).

Cupped invaginations filled cytoplasm with mitochondria, ribosomes and lipids are characteristic for many plastids of wheat callus cells. Similar invaginations are observed in leucoplasts of nectary cells of *Heracleum sp*. in the period of flowering maximum, in nectary cells of *Cucumis sativus* L. in the final stage of aging and immediately before dying away, in the cells of glands of *Ribes nigrum* L. secreting essential oil, and in the cells of needles of *Pinus sibirica* Du Tour. (Vassilyev, 1977). Described the structural features are also characteristic for plastids found in secretory cells of vascular plants, which produce secondary metabolites - glycoproteins and terpenes (Vassilyev, 2000; Kolalite, Ivanova, 2002).

In the initial stages of apoptosis of wheat callus cells plastids with prolamellar bodies, with the weak development of the membrane system and plastoglobules grouped around prolamellar body, and very rarely - with starch grains are generally marked in the cytoplasm height vacuolated cells. Many plastids are cupped. Such elongated plastids with invaginations are also found in cells with features of programmed death in coleoptiles wheat (Vanyushin, 2001). In the final stages of degradation of wheat callus cells, for our visual evaluation, the number of plastids decreases.

Plastids of barley callus cells having a different shape, actively accumulate starch. The stroma of the plastids is moderate electron dense and has well defined tubular peripheral reticulum, underdeveloped lamellar system and the low number of plastoglobules, which have higher electron density than the stroma. The presence of membrane system in organelles involves them in the process of photosynthesis. Apparently, part of the starch accumulated in plastids, is a product of photosynthesis, and the remaining part is produced from sucrose present in the culture medium (Galiyeva, 2001).

### Mitochondria

In wheat callus cells mitochondria are elongated (average of (1.4 x 0.4) μm), with long cristae and matrix, which is somewhat denser than the cytosol. The mitochondria have clearly visible mitoribosomes, intramitochondrial granules are observed rarely.

In the process of apoptosis mitochondria lose their shape, their matrix is enlightened, cristae become greatly shortened and swell. It should be noted that we have not found dense mitochondria with condensed mitochondrial matrix, characteristic for apoptotic cells of wheat (Bakeeva, 2003). We observed the patterns of mitochondrial degradation relevant mechanisms mitoptosis (Skulachev, 2001), in which the matrix of the organelle becomes swelling, folds of the inner membrane (cristae) – straighten out, and the outer membrane – breaks.

In barley callus cells chondriome is well developed; mitochondria are slightly elongated (average of (0.7 x 0.4) μm). Interestingly, mitochondria are retained for quite a long time until the last stages of cell degradation. They remain morphologically intact until rupture of the tonoplast. Similar morphological features occur during the premature cell death of anther tissues in sunflower (Balk and Leaver, 2001). These authors showed that the cells and nuclei condensed, and mitochondria persisted until extremely late in the cell death process. If given that the programmed cell death is energy dependent process, the long-term preservation of the mitochondria, which provide energy the cell before the last stages of death, is understandable. The death of the mitochondria requires no proteins other than those that are in the organelle (Skulachev, 2001). It is now an established fact that mitochondria not only generate energy for cellular activities but also play an important role in programmed cell death (Shah et al., 2000). Perhaps, organization of cytoplasm in the form of apoptotic bodies (membrane-bounded cytoplasmic fragments containing gradually degraded mitochondria and other organelles and inclusions) promotes long-term protection of the structure and activity of mitochondria in the process of physiological cell death (Bakeeva et al., 1999).

In addition, some authors (Reape et al., 2008) suggest that the death programme is regulated in complex ways and the mitochondria and more specifically the permeability of the outer mitochondrial membrane is possible regulator of apoptosis.

### Vacuole

Programmed cell death in wheat callus is manifested by a gradual decrease in the volume of the cytoplasm and a concomitant increase in the volume occupied by lytic vacuoles. The presence of large vacuoles is typical for degraded cells, cell component consists of a narrow layer located near the cell wall; there are also small vacuoles with bubbles in this place. Part of vacuoles contains sparse sediment, most of them has small bubbles. Invaginations in the vacuolar membrane (tonoplast) and fusion of vesicles with the vacuole, followed by uptake and degradation of portions of the cytoplasm in the vacuolar lumen are often observed in electron micrographs. Rupture of the tonoplast is the final step in the execution of programmed cell death in wheat callus. Apparently, tonoplast of vacuole disintegrates quickly and vacuolar hydrolases are evenly distributed throughout the cell (Van Doorn et al., 2011). These data are in accordance with findings providing evidence for the rupture of the large central vacuole triggering the plant PCD (Zheng et al., 2017).

Programmed cell death of barley begins to gain the degree of vacuolization, like wheat. Vacuoles contain both normal vesicles and cytoplasmic portions with organelles (apoptotic-like bodies). Formation in the vacuole of vesicles with mitochondria is also a characteristic feature of the apoptotic cells in aging coleoptile of etiolated wheat seedlings (Bakeeva et al., 1999; 2001). These apoptotic-like bodies are engulfed and recycled by neighboring cells; therefore complete elimination of the cell occurs (Palavan-Unsal et al., 2005). Sometimes in the vacuoles there are electron-dense globules reaching the size of (6.89 x 4.82) μm and a density close to the lipid. Further degradation of cells occurs differently. Contents of some cells completely disappear, only electronic transparent vacuoles with the remnants of plastids and the bubbles keep and localized along the cell walls. In other cells forms the central vacuole with rare fibrils.

As is known, intense vacuolization of cells with organization of huge vacuoles, formation in the cellular vacuole of vesicles with subcellular organelles is one of the specific features that is common for apoptotic plant cells and promotes cell autolysis, their autodigestion (Zamyatnina et al., 2002). Some authors have reported that there are no features of degradation in plastids and mitochondria observed in the vacuolar vesicles and believe that the condition for the organelle survival is more comfortable in vesicles than that in other cytoplasmic compartments of the apoptotic plant cell (Zamyatnina et al., 2002). In our view mitochondria are stored in apoptotic-like bodies for a long time, but start to degrade after rupture of the tonoplast.

### Fibrillar material

Electron microscopic study allowed us to detect the release of electron-dense substances in the periplasmic space and the surface of apoptotic cells (Bishimbayeva, 2002).

Thus, degrading cells of wheat and barley callus have all known ultrastructural features, which are characteristic for apoptotic cells: intense vacuolization, increasing the size of the nucleus, cupped invaginations of plastids, mitochondria saving until the last stages of cell degradation and the formation of vesicles, similar to apoptotic bodies.

In addition, we the first showed that degrading cells in embryogenic callus are characterized with the destruction of the nuclear membrane and the allocated substances of fibrillar nature in the periplasmic space and on the surface of cells.

